# Food availability early in life impacts among and within individual variation in behaviour

**DOI:** 10.1101/2023.02.23.529667

**Authors:** Cammy Beyts, Julien G. A. Martin, Nick Colegrave, Patrick Walsh

## Abstract

1. The availability of food during early life has been proposed as a key proximate mechanism for the development of variation in behaviour among and within individuals.
2. Individuals can vary amongst each other in their personality, plasticity and predictability and if an individual’s behaviour is correlated across contexts this can lead to behavioural, plasticity and predictability syndromes.
3. In this study, we used a split brood design to raise African clawed frog tadpoles (*Xenopus laevis*) on a high or low diet in food availability and measured the distance they swam in a familiar and unfamiliar context eight times during their development.
4. In a familiar context, we found that there was an increase in among individual variance in plasticity and predictability in the high food treatment. This shows that when resources are not restricted, individuals are not constrained in the expression of their behaviour at certain phenotypic levels.
5. In an unfamiliar context, we found a different response, with an increase in individual variance in personality in the low but not the high feed tadpoles. As unfamiliar contexts may be riskier, our results highlight that individuals receiving less food may take greater foraging risks in novel contexts.
6. Across contexts, we found a predictability syndrome in the high but not the low feed tadpoles, highlighting that cross-context behaviours can become decoupled in some developmental conditions but remain intact in others.
7. Together our findings show that early life conditions contribute to among individual variation in behaviour but that these may only impact the phenotype at specific phenotypic levels and are context specific.
8. We emphasise that having a fundamental understanding of how early development may promote or constrain individual variation can provide a greater understanding of how individuals and populations may respond to novel conditions brought about by anthropogenic activity.

## Introduction

A major goal in behavioural ecology is to understand why individuals from the same population differ from each other in their behavioural responses (Sih et al. 2004; Réale et al. 2007; Réale et al. 2010; Dingemanse et al. 2012). This among individual variation is in part due to genetic differences between individuals but also to how different genotypes respond to differences in the environment, a process known as behavioural plasticity (Snell-Rood 2013). Behavioural plasticity can occur early in life or later in development, processes known as developmental plasticity or environmental plasticity respectively (Wilson et al. 2010; Dingemanse and Wolf 2013; Snell-Rood 2013). During early development, individuals are particularly sensitive to environmental conditions such as food availability (Han and Dingemanse 2015; Fischer et al. 2016; Stamps 2016; Urszán et al. 2018), which can have long lasting impact on a large diversity of traits from morphology, physiolgy and behaviours. This can mean that two individuals who develop under high or low food resource conditions may show very different behavioural responses to the same ecological context later in life, despite being from a similar genetic background (Han and Dingemanse 2015; Han and Dingemanse 2017; Royauté and Dochtermann 2017; Royauté et al. 2019; Fanson et al. 2021). Understanding how early-life conditions impact among individual variation in behaviour is of growing interest because it can explain how individuals from different parts of a specie’s range can maintain high fitness, despite being exposed to very different environments (Dingemanse and Wolf 2013; Urszán et al. 2015; Stamps 2016; Urszán et al. 2018). Populations harbouring higher rates of among individual variation also suggest a greater potential for individuals to display behavioural responses which may be adaptive if the environment changes as a result of anthropogenic disturbance (e.g. urbanisation and climate change) (Chevin et al. 2010; Dingemanse and Wolf 2013; Sol et al. 2013). Therefore, having a fundamental understanding of how the early environment may promote or constrain among individual variation could provide a greater understanding of how individuals and populations may respond to novel environments.

During development animals are often restricted in their movements or constrained to certain habitats, making them dependent on the local food resources in these locations (Han and Dingemanse 2015). As the amount of food available is rarely uniform across a landscape, food availability during development has been highlighted as a key resource that has the potential to influence the phenotypic expression of behavioural responses both within and across populations (Han and Dingemanse 2015; Han and Dingemanse 2017). However, the impact of food availability on behavioural responses can be difficult to anticipate because individuals can vary amongst each other at multiple phenotypic levels and how an individual behaves may also depend on the ecological context that a behavioural response is measured (Westneat et al. 2015; Mitchell and Houslay 2021; O’Dea et al. 2021). Consequently, it is necessary to understand how individuals may vary in their behaviour at different phenotypic levels and across multiple contexts to fully understand how the phenotype responds to early developmental conditions.

There are three key aspects to behavioural variation: personality, plasticity and predictability. The behavioural response of an individual at one point in time can be quantified as their initial behaviour (Réale et al. 2007; Kelleher et al. 2018). If the same individual is measured on repeated occasions, it is also possible to determine how their behavioural response changes over time (Dingemanse et al. 2002; Martin and Réale 2008; Dingemanse et al. 2012; Stamps and Biro 2016). The former can be described as an animal’s personality and the latter as its behavioural plasticity (Dingemanse et al. 2010; Martin et al. 2011; Stamps 2016). For example, the distance an individual travels within their familiar home territory could be described as their initial foraging activity or personality. Whereas the change in an individual’s activity level when measured one week later would be described as their plasticity. After accounting for personality and plasticity, an individual may further show short-term changes in their behavioural response, known as behavioural predictability (Stamps et al. 2012; Cleasby et al. 2015a; Westneat et al. 2015; Mitchell et al. 2016). Unpredictable individuals will show inconsistencies in their behavioural response, whereas predictable individuals will show more consistent behavioural responses when measured over a short time span (Stamps et al. 2012; Westneat et al. 2015) (Figure 1).

**Figure 1.**
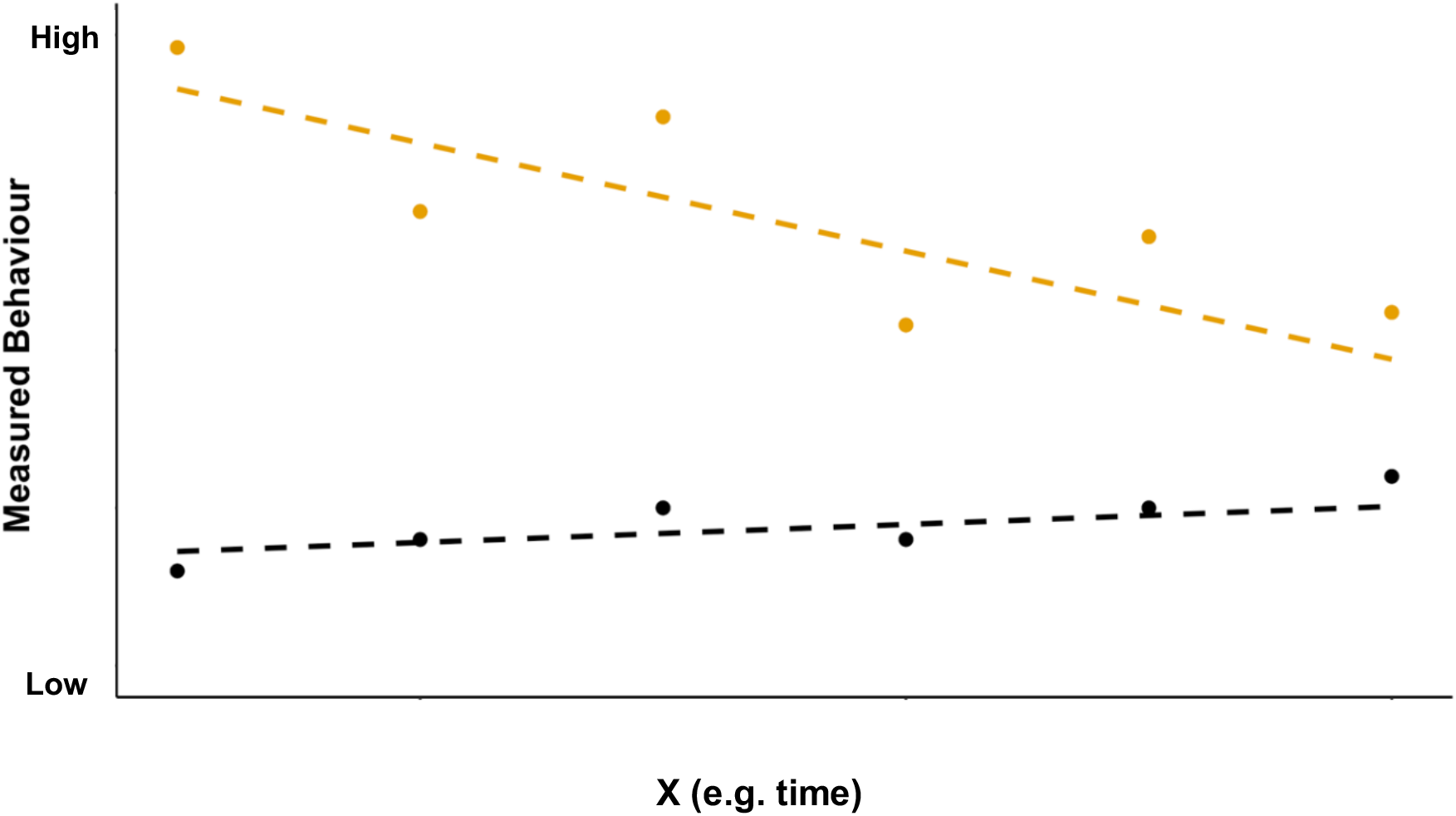
Conceptual diagram of two individuals who differ in their initial behaviour (personality), behavioural plasticity and behavioural predictability. The orange individual has a greater behavioural response initially (higher intercept) and a greater level of behavioural plasticity (steep slope) compared to the black individual. The points around the orange individual’s slope are also wider, indicating that the orange individual had a lower level of predictability (higher residual variance) compared to the black individual. Figure adapted from (Stamps et al. 2012; Jolles et al. 2019).

Where there is a correlation between an individual’s personality, plasticity and predictability across contexts, these are known as behavioural, plasticity and predictability syndromes respectively (Houslay et al. 2018; Mitchell et al. 2021a; Mitchell and Houslay 2021; O’Dea et al. 2022).

The amount of food available during development may influence among individual variance in personality, plasticity and predictability by impacting the internal state of individuals such as their growth rate, metabolism or energy reserves (Han and Dingemanse 2015; Royauté and Dochtermann 2017; Royauté et al. 2019). When developing under high food conditions, natural genetic variation in metabolic rate within a population may be unconstrained, resulting in an increase in among individual variance in personality, plasticity and/or predictability (Hoffmann and Merilä 1999; Charmantier and Garant 2005; Dingemanse and Wolf 2010; Dingemanse and Wolf 2013; Snell-Rood 2013). For example, individuals with faster metabolic rates may show higher initial foraging activity, sustain high levels of foraging activity over their lifespan and be more consistent in their foraging activity over time to meet their metabolic requirements. In contrast, individuals with slower metabolic rates may show lower initial foraging activity and due to their lower metabolic needs, may be more plastic or less predictable in their foraging behaviour (Careau et al. 2008; Biro and Stamps 2010; Biro et al. 2018). However, when developing under low food availability, individuals with faster metabolic rates may be constrained, resulting in all individuals developing a similar slow metabolic phenotype and reducing among individual variance in personality, plasticity and/or predictability (Hoffmann and Merilä 1999; Charmantier and Garant 2005; Wolf et al. 2008; Snell-Rood 2013).

The impacts of food availability on among individual variance in personality, plasticity and predictability may also be context dependent (Stamps and Groothuis 2010; Han and Dingemanse 2015; Mitchell and Houslay 2021). In an unfamiliar context, foraging may be more metabolically expensive compared to a familiar context as individuals need to remain vigilant against potential predator threats (Houslay et al. 2018; Mathot et al. 2019). For example, when developing under low food availability, the increased need for food may highlight variance among individuals in their propensity to take risks, increasing among individual variance in behaviour at one or multiple phenotypic levels relative to a familiar context (McNamara and Houston 1994; Han and Dingemanse 2017; Royauté and Dochtermann 2017; Royauté et al. 2019; Fanson et al. 2021). Food availability may also impact how individual’s behaviour is correlated across different ecological contexts (Endler 1995; Han and Dingemanse 2015; Sikkink et al. 2015). Under low food availability, individuals may have to forage more intensely across a range of contexts to secure sufficient resources, resulting in behavioural, plasticity and/or predictability syndromes (Bell and Sih 2007; Dingemanse et al. 2007). In contrast, adequate resources may mean that individuals have the ability to adjust their behaviour according to the ecological context they are in, resulting in the absence of behavioural, plasticity and/or predictability syndromes.

Anuran larvae are an excellent group of organisms to investigate how early development under high or low food availability may impact among individual variance in behaviour. (Relyea 2001; Martin and Pfennig 2012; Levis et al. 2015; Urszán et al. 2015; Urszán et al. 2018). In species such as *Xenopus leavis*, adults produce large number of eggs in a single egg mass, which is advantageous for split brood designs, where full siblings can be reared under different early developmental environments. This limits the genetic differences between experimental groups, making proximate effects on behaviour less ambiguous (Stamps 2016). Large numbers of individuals per egg mass also allow for higher sample sizes required for estimating among individual differences in behavioural plasticity and predictability (Martin et al. 2011; Cleasby et al. 2015b). As tadpoles are also geared towards growth, this makes them highly responsive to resource availability during development (Martin and Pfennig 2012; Wilson and Krause 2012a; Wilson and Krause 2012b).

In this study, we investigated how the availability of food during development impacts among individual variation in the distance *X. leavis* tadpole swam in familiar and novel contexts. In each context, we were interested in how food availability impacted 1) among individual variance in the initial distance tadpoles swam (personality), 2) among individual variance in how tadpoles adjusted the distance they swam over a series of 8 trials (plasticity) and 3) among individual variance in the consistency of distances swam trial to trial (predictability). We were also interested in how food availability impacted tadpole behaviour across contexts by examining if there was a correlation between the 4) initial distance tadpoles swam in both contexts (behavioural syndrome), 5) how tadpoles adjusted the distance they swam from trial 1 to 8 across contexts (plasticity syndrome) and 6) consistency in the distance tadpoles swam trial to trial in each context (predictability syndrome).

## Methods

### Husbandry and Feeding protocols

We obtained seven egg masses of wild type *X. laevis (Xenopus* Resource Centre, Portsmouth, UK). The egg masses were reared separately, in batches of 200 embryos, in 300 x 195 x 205 mm growth tanks (Exo Terra, UK). Embryos from the same egg mass were reared in the same tank so we could identify which egg mass embryos/tadpoles originated from. Tanks were filled with 4 litres of treated (Seachem Prime, USA) and aerated tap water to remove chlorine and chloramine. Water temperature was maintained at 21°C throughout the experiment with a 12-hour light/dark cycle. Tadpoles were fed 0.05g of Sera Micron (Sera, Germany) five days per week over 10 days. After 10 days we randomly assigned tadpoles in each growth tank onto one of two feeding treatments: high feed or low feed treatments.

To account for differences in the number of tadpoles across the stock tanks (mean number of tadpoles per tank = 54.4 ± 8.8 SD), all tadpoles were raised under the same tadpole density (0.08 litres of water per tadpole) and tadpoles were fed according to the number of individuals within a tank. Individuals in the high feed treatment were fed 0.0015g of Sera Micron per tadpole and individuals in the low feed treatment were fed 0.0005g of Sera Micron per tadpole. To account for growth increases, the feed was increased to 0.003g and 0.0015g of Sera Micron for the high and low feed tadpoles respectively after 10 days. Tadpoles were fed 5 days per week (Monday-Friday).

Seven days prior to starting behavioural trails, we randomly selected tadpoles from across the stock tanks and transferred 36 tadpoles from each of the high and low feeding treatments into one of 72 “home” tanks measuring 170 x 130 x 130 mm (Insta, China). Tanks were filled with 1 litre of water and housed one tadpole per tank to allow individual identification throughout the rest of the study. Tank sides facing other tadpoles were obscured to ensure tadpoles did not receive visual cues from other tadpoles.

The seven-day acclimatisation period was to allow tadpoles to adjust to their new tank conditions ahead of starting behavioural trials. This was important as tadpole behaviour was filmed in an individual’s “home tank” to measure the distance tadpoles swam in a familiar context. The tanks were part of a closed recirculating water system in which the water within each of the 72 x 1 litre tanks was continually replaced over a ~ 30-minute period. Water exiting the tanks was passed through physical filters to remove food and tadpole excretion products and biological filtration was used to control ammonia and nitrite levels. As small tadpoles risked passing through the tank barriers and into the water filtration system, tadpoles that were below Xenopus stage 48 (Nieuwkoop and Faber 1994) when randomly selected were returned to their stock tanks and new individuals were selected. We continued to feed tadpoles five times per week and due to the continuous water flow in the tanks, feeding was increased to 0.3g per tadpole in the high feed treatment and 0.1g per tadpole in the low feed treatment. Water flow was paused for 45 minutes when adding food to the home tanks to allow tadpoles time to feed. Tadpoles were fed between 5-7pm each day to fall after the completion of the day’s behaviour trials and to ensure tadpoles were fed on the evening prior to behavioural trials on the following day.

Behavioural trials were filmed over two experimental “sets”, with the second set of tadpoles being placed into home tanks (cleaned with aquarium disinfectant) after tadpoles from the first set had completed all of their behavioural assays. Our final data set contained data from 60 individuals in the high food treatment and 42 individuals in the low feed treatment.

### Behaviour assays

We recorded the total distance each tadpole swam in a familiar context and an unfamiliar context in two separate behavioural assays, commonly referred to as activity and exploration behavioural responses in the behavioural ecology literature (Réale et al. 2007; Kelleher et al. 2018). The distance each tadpole swam in each context was recorded on eight separate occasions to give a total of 960 familiar and unfamiliar context trials in the high food treatment and 672 familiar and unfamiliar context trials in the low feed treatment. We were blind to the treatment identity of tadpoles during all assays. There were 17 tadpole deaths whilst filming behavioural trials (10 high feed, 7 low feed). Partial recordings from tadpoles which did not complete all 8 trials were removed. For each trial, the distance travelled in a familiar and unfamiliar context was recorded on the same day, with the least disruptive familiar context assay recorded first and the unfamiliar context assay recorded immediately after, to limit the carry over effects of any stress which may have been induced by the unfamiliar context (Bell 2013). All eight trials were conducted within a 14-day period and there was an average duration of 1.25 days between an individual’s consecutive familiar context trials.

The distance tadpoles travelled in the familiar and unfamiliar contexts was recorded using four Canon Legria HF R86 camcorders, which were fixed in positioned 450 mm above the tadpole’s home tank (in the case of a familiar context) or unfamiliar tank. Two tadpoles in separate, adjacent tanks could be filmed simultaneously under one camera. All tank sides were obscured to prevent social cues from other tadpoles or accidental disturbance from the experimenter during filming. The familiar and unfamiliar tanks could be positioned and removed from under the camera but were held in a fixed position during trials to assist with automotive tracking software. During trials, the tanks were lit from underneath using an A3 sized light box to provide an equal level of lighting across the arena, preventing shadows from forming. The distance tadpoles swam in familiar and unfamiliar contexts were filmed in a recording room, adjacent to the lab where husbandry procedures took place so that tadpoles would be undisturbed during filming. The recording room had the same temperature and lighting conditions as the husbandry room. Post filming, all videos were re-sized to 640×360 pixels and reduced from a frame rate of 25fps to 1fps using the command line tool ffmpeg (Tomar 2006). In both contexts, we measured the total distance a tadpole travelled in pixels using a custom-written tracking tool (written by CB, see data availability) developed in Python v3.0 and using the OpenCV v4.4 library.

### Familiar context

To measure the total distance tadpoles swam in a familiar context, we recorded the movement of tadpoles in their home tanks over a 10-minute period. Prior to filming, all tadpoles were left undisturbed for 10 minutes to acclimatise to the recording room environment.

### Unfamiliar context

To measure the total distance tadpoles swam in a novel context, we recorded the movement of tadpoles in a tank they were unfamiliar with. The unfamiliar tank was larger than their home tank and measured 203mm x 203mm (iDesign, UK). Before tadpoles were added to the tank, it was filled with 500ml of treated and aerated 21°C tap water. To start a trial, tadpoles were transferred from their home tank to the novel tank using a new 7ml bijou container. Tadpole movements were recorded over a 20-minute period. Only the first 10 minutes of footage was used in the analysis so that the distance travelled in the unfamiliar context could be more easily compared to the familiar context. The tank was cleaned between trials using tap water and fresh water was used for each new trial and tadpole.

### Morphological measures

The snout vent length (SVL), a measure of body size, of each tadpole was measured in FIJI v2.0 (Schindelin et al. 2012) to the nearest 0.1mm from recordings taken from the unfamiliar context. Measurements were taken from each unfamiliar context trial to give eight SVL measurements for each tadpole.

### Ethical approval

All procedures were approved by the University of Edinburgh ethics committee, under the assessment pwalsh1-0001.

### Statistical analysis

To estimate the effect of feeding treatment i) the variance among individuals in the distance they swam in their first trial (personality), ii) the variance among individuals in the change in distance they swam from trial 1 to 8 (plasticity), and iii) the variance among individuals in the consistency of the distance they swam trial to trial (predictability) in both a familiar and unfamilar context we used a bivariate double hierarchical generalized linear model (DHGLM) (Lee and Nelder 2006; Cleasby et al. 2015b). The bivariate DHGLM allowed all parameters i-iii to be estimated for both the total distance swam in a familiar and unfamiliar context simultaneously as well as their pairwise correlations to estimate iv) behavioural, v) plasticity and vi) predictability syndromes. The mean part of the model was used to estimate treatment effects on among individual variance in personality and plasticity, while the dispersion part of the model was used to estimate among individual variation of behavioural predictably. The mean and dispersion model also allowed us to model how feeding treatment impacted the vii) average distance tadpoles swam and the viii) average predictability of tadpole swimming behaviour at the population level in each context.

We fitted the same model structure for both behavioural responses, the distance tadpoles swam in a familiar and an unfamiliar context. To estimate the effect of feeding treatment on the average distance tadpole swam, we included a fixed effect of treatment (high vs low) in the mean model. To estimate how tadpoles changed the distance they swam from trial 1 to 8, i.e. habituation, we included a fixed effect of trail number, fitted as a continuous covariate with the first trial coded as zero. In addition, we included tadpole body size (SVL scaled and mean centred) and experimental set (set 1 and 2) to control for changes in tadpole body size (SVL) and potential differences due to experimental conditions in the two sets. To estimate among individual variance in initial behaviour and plasticity in habituation, we fitted individual identity as a random intercept and random slope term with trial number in the mean model. To estimate the effect of feeding treatment on among individual variance in initial behaviour and plasticity, we estimated treatment specific random effects for both random intercept and random slope terms similar to a character state approach. For the dispersion part of the model, the model included feeding treatment as a fixed effect and treatment specific random effects for individual identity.

Estimating among individual variance in personality, plasticity and predictability in the distance travelled in familiar and unfamiliar contexts for high and low feeding treatments created two 6×6 covariance matrixes, one matrix for each treatment. The trait covariance matrices allowed us to estimate covariance in initial behaviour, behavioural plasticity and behavioural predictability within contexts and behavioural, plasticity and predictability syndromes between contexts. Given that tadpoles were only exposed to one feeding treatment, we could not estimate the covariance across treatments at the individual level. Finally, given that we measured the distance tadpoles swam in a familiar context and then measured the distance tadpoles swam in an unfamiliar context straight after, we also estimated the within individual (residual) correlation between contexts in the high and low food treatments. We included a random term for egg mass identity in the mean and dispersion models to control for variation associated with egg mass identity. However, this term was subsequently removed as there was minimal variation in tadpole behaviour associated with egg mass identity and only a small number of egg masses (n = 7). The fit of the models with and without egg mass identity were similar when compared using leave-one-out cross-validation (LOOIC) using the loo package v4.2 in R (Vehtari et al. 2017).

For individual *i* at observation *t*, the bivariate DHGLM can be written as

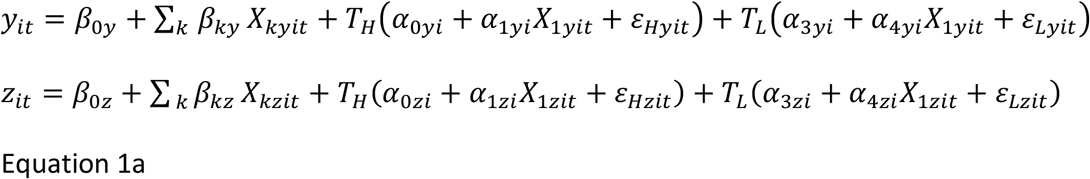

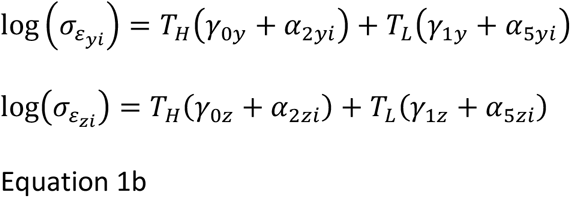

Where *y* and *z* are the two behavioural responses, distance travelled in a familiar and unfamiliar context. Subscripts_*y*_ and _*z*_ indicate context specific estimates and variables. *β* and *γ* represent fixed effect parameters in the mean and dispersion part of the model respectively. *X* are fixed effects with *X*_1_ being the observation number. *α* are the individual deviations from the average individual for the intercept (0 and 3), plasticity (1 and 4) and predictability (2 and 5) estimates. *T_L_* and *T_H_* are dummy variables coded as zero/one for low and high feeding treatments allowing to estimate treatment specific variance parameters for the random effects and the dispersion (equation 1a–b). Treatment specific random effects (_o–2_ for high food and _3–5_) have a multivariate normal distribution (equation 2a–b)

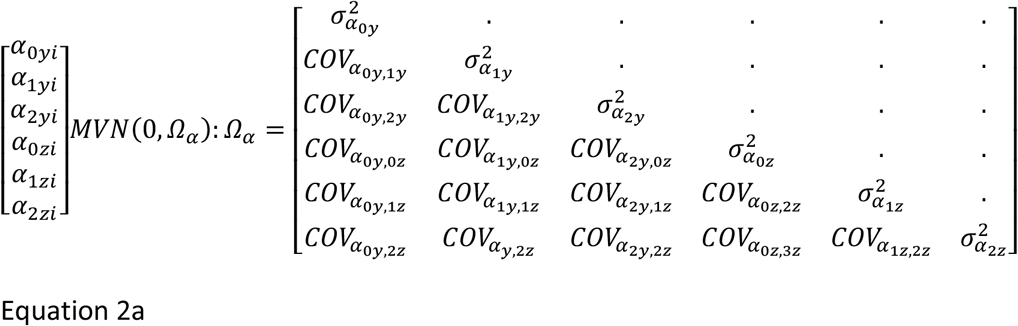

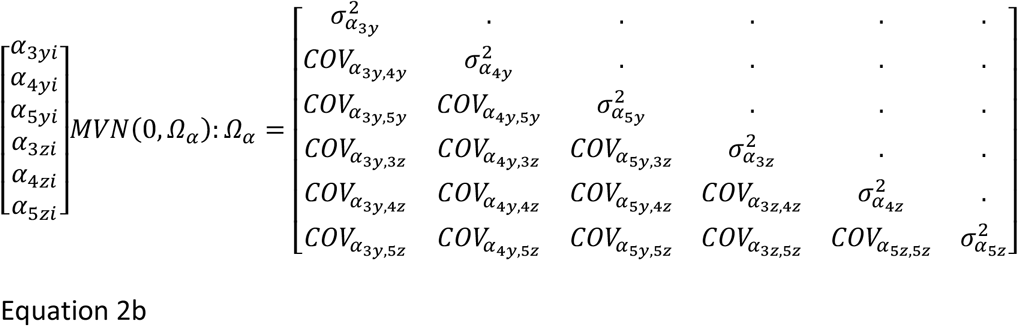

For each treatment, the residuals follow a multivariate normal distribution (equation 3a–b) where *r_ε_* is the residual correlation between the behavioural responses in a familiar and unfamiliar context.

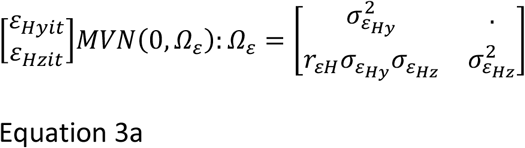

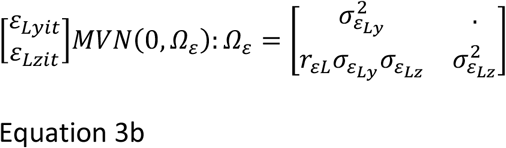

The bivariate DHGLM was fitted in a Bayesian framework using stan v2.21.0 (Stan Devlopment Team. 2020) fitted with the rstan package v 2.21.2 (Stan Devlopment Team. 2020a) within R 4.0.0 (RCoreTeam. 2013). We used uninformative or weak priors on all parameters (Gelman et al. 2013) which included wide normal priors for fixed effects, half-student priors for variance parameters and LKJ correlation priors for correlations. For all parameters, autocorrelation was below 0.1, all chains had converged (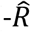 below 1.1) and effective sample size was large (Gelman et al. 2013). Posterior predictive checks showed an adequate fit of the model

To establish whether the effect of treatment resulted in a change in the estimated parameters for among individual variance and correlations, the difference in the variance/correlation parameter between the high minus the low feeding treatment was calculated at each iteration, obtaining the posterior distribution of the difference. For all parameters, we then reported the mean of the posterior distribution with the highest posterior density interval (HPDI) at 95%.

## Results

In a familiar context, tadpoles in the high feed treatment swam shorter distances on average compared to tadpoles in the low feed treatment (Figure 2a, Table 1). There was no difference in the average distance swam between food treatments in an unfamiliar context (Figure 2b, Table 1).

**Figure 2.**
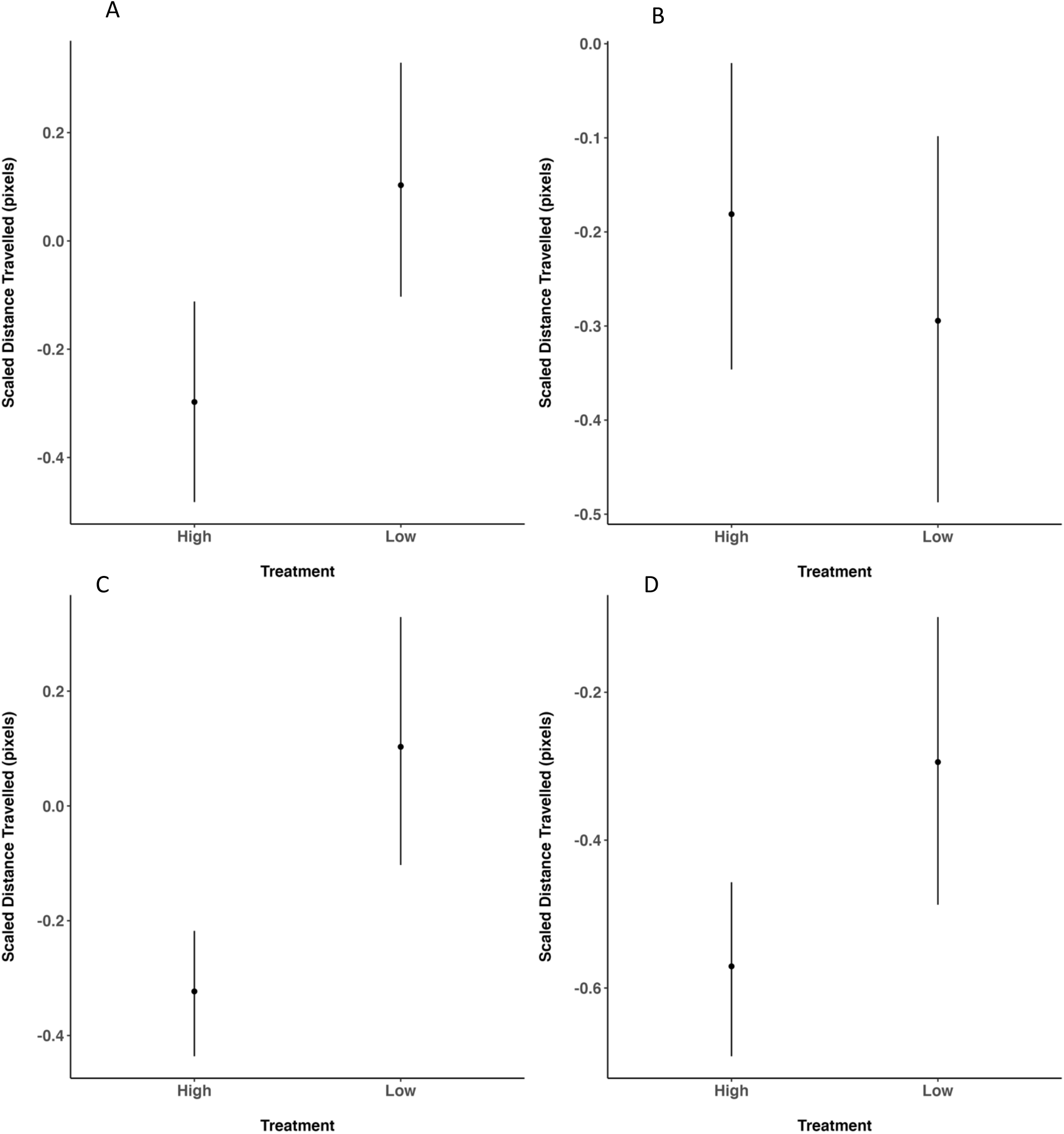
Posterior mean estimate and 95% credible intervals for the effect of feeding treatment on overall distance travelled in a familiar (A) and unfamiliar context (B) as well as the overall predictability of distance travelled in a familiar (C) and unfamiliar context (D).

**Table 1.** Posterior estimates for the effect of feeding treatment and the treatment comparison for the average distance swam, the average predictability, as well as the variance among individuals in the distance tadpoles swam in their first trial (personality), change in distance swam from trials 1 to 8 (plasticity) and the predictability of distance swam in a familiar and unfamiliar context. Also displayed are the population estimates for body size (SVL), trial and set. Estimates are displayed alongside their 95% credible intervals (CI).

|  | Familiar Context |  |  | Unfamiliar Context |  |  |
| --- | --- | --- | --- | --- | --- | --- |
|  | Mean | 95% CI |  | Mean | 95% CI |  |
|  |  | 2.5 | 97.5 |  | 2.5 | 97.5 |
| Fixed effects |  |  |  |  |  |  |
| SVL | 0.087 | -0.006 | 0.177 | 0.302 | 0.228 | 0.383 |
| Trial | 0.066 | 0.033 | 0.097 | 0.072 | 0.040 | 0.105 |
| Set | -0.194 | -0.395 | -0.002 | -0.085 | -0.265 | 0.094 |
| Population: Mean behaviour |  |  |  |  |  |  |
| High | -0.297 | -0.482 | -0.112 | -0.181 | -0.346 | -0.020 |
| Low | 0.103 | -0.103 | 0.329 | -0.294 | -0.487 | -0.098 |
| High - low | -0.400 | -0.588 | -0.204 | 0.113 | -0.066 | 0.298 |
| Population: Predictability |  |  |  |  |  |  |
| High | -0.323 | -0.436 | -0.218 | -0.571 | -0.692 | -0.457 |
| Low | -0.066 | -0.158 | 0.028 | -0.605 | -0.749 | -0.475 |
| High - low | -0.257 | -0.409 | -0.130 | 0.034 | -0.135 | 0.207 |
| Variance among individuals: Personality |  |  |  |  |  |  |
| High | 0.044 | 0.000 | 0.102 | 0.042 | 0.000 | 0.097 |
| Low | 0.114 | 0.000 | 0.249 | 0.165 | 0.058 | 0.289 |
| High - low | -0.070 | -0.226 | 0.075 | -0.123 | -0.259 | 0.001 |
| Variance among individuals: Plasticity |  |  |  |  |  |  |
| High | 0.011 | 0.003 | 0.019 | 0.019 | 0.010 | 0.029 |
| Low | 0.003 | 0.000 | 0.008 | 0.013 | 0.005 | 0.022 |
| High - low | 0.008 | -0.002 | 0.017 | 0.006 | -0.010 | 0.018 |
| Variance among individuals: Predictability |  |  |  |  |  |  |
| High | 0.106 | 0.042 | 0.168 | 0.120 | 0.044 | 0.199 |
| Low | 0.021 | 0.000 | 0.057 | 0.127 | 0.047 | 0.224 |
| High - low | 0.086 | 0.010 | 0.160 | -0.006 | -0.131 | 0.117 |

Considering the distance tadpoles swam in their first trial, there was no evidence of among individual variation in the initial distance swam in a familiar context in either treatment, whereas for an unfamiliar context, there was among individual variation in the low feed treatment but not the high feed treatment (Table 1). Turning to plasticity, in a familiar context, tadpoles in the high feed but not the low feed treatment showed among individual variance in plasticity (Table 1). In the unfamiliar context, tadpoles across both treatments showed among individual variance in plasticity which was not found to be larger in one treatment than the other (Table 1).

Tadpoles in the high feed treatment were on average more predictable in the distances they swam compared to tadpoles in the low feed treatment in both a familiar and unfamiliar context (Figure 2c and Figure 2d, Table 1). However, whilst high feed treatment tadpoles were on average more predictable, there was also more among individual variation in tadpole predictability in the high feed treatment compared to the low feed treatment in a familiar context (Table 1). In the unfamiliar context, there was also a mix of predictable and non-predictable tadpoles across both treatments, but among individual variance in behavioural predictability did not differ between treatments (Table 1).

Apart from low feed tadpoles in a familiar environment, tadpoles which showed greater plasticity in their swimming responses from trial 1 to 8 were also less predictable in the distances they swam trial to trial, resulting in a positive correlation between individual estimates of plasticity and predictability (Table 2). This positive association between plasticity and predictability was consistent between treatments in both contexts (Table 2). Within the low feed treatment, we found that tadpoles which swam further in their first trial also showed a steeper change in the distance they swam between their first and last trial, leading to a positive correlation between individual estimates of personality and plasticity (Table 2). We did not find that this correlation between personality and plasticity changed between treatments in either context (Table 2).

**Table 2.** Posterior correlation estimates for among individual and within individual correlations and 95% credible intervals (CI), for high and low feed tadpole groups as well as the difference between feeding treatment groups. Names of the parameters starting with “FC” and “UFC” refer to familiar context and unfamiliar contexts respectively. Estimates with CI which do not overlap zero have been highlighted in bold.

| Correlation Parameter | High Feed |  |  | Low Feed |  |  | High – Low Feed |  |  |
| --- | --- | --- | --- | --- | --- | --- | --- | --- | --- |
|  | mean | 95% CI |  | mean | 95% CI |  | mean | 95% CI |  |
|  |  | 2.5 | 97.5 |  | 2.5 | 97.5 |  | 2.5 | 97.5 |
| Among individuals |  |  |  |  |  |  |  |  |  |
| Behavioural syndrome | 0.416 | -0.298 | 0.846 | 0.018 | -0.524 | 0.553 | 0.398 | -0.444 | 1.190 |
| Plasticity syndrome | 0.023 | -0.353 | 0.384 | 0.139 | -0.566 | 0.723 | -0.116 | -0.894 | 0.614 |
| Predictability syndrome | **0.452** | **0.102** | **0.741** | 0.337 | -0.358 | 0.823 | 0.114 | -0.489 | 0.829 |
| Personality vs plasticity |  |  |  |  |  |  |  |  |  |
| FC Personality FC Plasticity | 0.160 | -0.431 | 0.720 | 0.037 | -0.630 | 0.704 | 0.123 | -0.755 | 1.022 |
| UFC Personality UFC Plasticity | 0.106 | -0.423 | 0.650 | 0.112 | -0.352 | 0.611 | -0.006 | -0.703 | 0.743 |
| UFC Plasticity FC Personality | 0.196 | -0.374 | 0.706 | 0.000 | -0.545 | 0.511 | 0.196 | -0.595 | 0.893 |
| UFC Personality FC Plasticity | 0.064 | -0.484 | 0.608 | 0.085 | -0.588 | 0.711 | -0.021 | -0.812 | 0.824 |
| Personality vs predictability |  |  |  |  |  |  |  |  |  |
| FC Personality FC Predictability | 0.516 | -0.067 | 0.887 | 0.308 | -0.488 | 0.839 | 0.208 | -0.587 | 1.114 |
| UFC Personality UFC Predictability | 0.403 | -0.179 | 0.846 | **0.495** | **0.049** | **0.832** | -0.093 | -0.742 | 0.589 |
| UFC Personality UFC Predictability | 0.337 | -0.166 | 0.780 | 0.130 | -0.489 | 0.692 | 0.207 | -0.560 | -0.917 |
| UFC Predictability FC Personality | 0.342 | -0.259 | 0.800 | 0.157 | -0.429 | 0.657 | 0.185 | -0.515 | 0.986 |
| Plasticity vs predictability |  |  |  |  |  |  |  |  |  |
| FC Plasticity FC Predictability | **0.691** | **0.350** | **0.920** | 0.211 | -0.583 | 0.814 | 0.479 | -0.222 | 1.253 |
| UFC Plasticity UFC Predictability | **0.561** | **0.226** | **0.825** | **0.479** | **0.071** | **0.809** | 0.082 | -0.406 | 0.582 |
| UFC Predictability UFC Plasticity | 0.112 | -0.278 | 0.489 | 0.146 | -0.521 | 0.747 | -0.034 | -0.749 | 0.717 |
| UFC Plasticity FC Predictability | 0.145 | -0.233 | 0.473 | 0.288 | -0.364 | 0.776 | -0.143 | -0.833 | 0.514 |
| Within individuals |  |  |  |  |  |  |  |  |  |
| Residual correlation | 0.027 | -0.073 | 0.131 | **-0.159** | **-0.279** | **-0.042** | **0.186** | **0.037** | **0.357** |

Turning to correlations between contexts, we found that high feed tadpoles which were more unpredictable in a familiar context were also more unpredictable in an unfamiliar context, resulting in a predictability syndrome (Table 2). However, we did not find a predictability syndrome in low feed tadpoles (Table 2). At the within individual level, we found that tadpoles which swam further distances in a familiar context, tended to swim a shorter distance in a unfamiliar context in the low feed but not the high feed treatment (Table 2). This resulted in a residual correlation across contexts which differed between the two treatment groups (Table 2).

## Discussion

We found that the amount of food available during development acted on multiple sources of among and within individual variation, with the largest impact on population and among individual estimates of behavioural predictability. In a familiar context, we found that high feed tadpoles swam further distances and were more predictable in the distances they swam on average. However, there was also a higher among individual variation in tadpole predictability in the high compared to the low feed treatment group. In an unfamiliar context, whilst high feed tadpoles were on average more predictable, we did not observe the same increase in among individual variance in behavioural predictability as seen in a familiar context. Together our results show that the amount of food available during development impacts population and among individual variance in behaviour at specific phenotypic levels which is also dependent on the ecological context an individual is measured in.

At a population level, we found that tadpoles in the high feed treatment swam shorter differences on average compared to tadpoles in the low feed treatment. This result seems contradictory since it might be expected that tadpoles in the low feed regime would have less energy and hence be less active than well fed tadpoles. A similar increase in the distance travelled under low-energy diets has been found in rodent species which can be attributed to a decrease in basal metabolic rate in order to allow more resources to be allocated to foraging activity (Overton and Williams 2004; Peña-Villalobos et al. 2020). As *X. leavis* are filter feeders, the lower density of food in the low feed treatment may have meant that tadpoles would have had to increase their buccal pumping rate and swim further distances to increase their feed intake (Wassersung and Hoff 1979; Ryerson and Deban 2010). Tadpoles have also been found to increase their gut length in response to food deprivation (Relyea and Auld 2004), which may form part of a wider strategy focused on more intense foraging efforts to compensate for nutritional deficiencies in their diet (Han and Dingemanse 2015)

The impact of food availability on among individual variance in behaviour has been mixed in previous studies. In response to food restriction, some studies have found that among individual variance in behaviour may increase while others have found this to decrease (Hoffmann and Merilä 1999; Charmantier and Garant 2005; Han and Dingemanse 2017; Royauté and Dochtermann 2017; Royauté et al. 2019). The results of this study reflect the diversity of these findings, as we found that the impact of food availability may be restricted to certain phenotypic levels or may change between ecological contexts. In a familiar context, our finding that among individual variance in predictability increased may reflect that only certain parts of an individual phenotype are developmentally plastic with respect to changing resources. For example, metabolic scope, the energetic limits within which an individual can draw from, has been highlighted as a distinct component of an individual’s metabolism and can lead to changes in an individual’s predictability in the absence of changes to their personality (Biro et al. 2018). Furthermore, we found evidence for among individual variance in personality and plasticity in the high but not the low feed treatment group. This supports the hypothesis that when resources are not restricted, individuals are not constrained in the expression of their phenotype, increasing among individual variance in behaviour (Sgrò and Hoffmann 2004; Charmantier and Garant 2005). However, in an unfamiliar context, it was low and not high feed treatment tadpoles which displayed among individual variance in personality. This finding supports the alternative hypothesis that among individual variance in behaviour will increase in response to food restriction (Hoffmann and Merilä 1999). This may be due to individuals differing in their ability to tolerate food restriction, causing low state individuals to take greater foraging risks in a novel context compared to higher state individuals (Dall and Griffith 2014). Consequently, to have a full understanding of the effects early development conditions have on the development of phenotypes, it is necessary to investigate the behaviour of individuals at multiple phenotypic levels and across distinct ecological contexts.

In the low feed treatment, tadpoles which were more exploratory in their first trial were also more unpredictable in the distance they swam in an unfamiliar context over time. However, the same response was not observed in tadpoles in the high feed treatment. Behavioural unpredictability has been hypothesised as a mechanism to reduce an individual’s risk of predation (Maye et al. 2007; Stamps 2007; Biro and Adriaenssens 2013; Briffa 2013; Richardson et al. 2018). Consequently, low feed tadpoles may have been highly motivated to explore due to a lack of food but owing to an increased risk of predation associated with higher exploration levels, may have increased their behavioural unpredictability to offset potential increased predation risks (Briffa 2013; Briffa et al. 2013; Horváth et al. 2019). Similar patterns have been found in Trinidadian guppies (*Poecilia reticulata*) where individuals with more ‘flight’-type stress responses in novel arenas were also less predictable (Prentice et al. 2020).

Except for low feed tadpoles in a familiar context, tadpoles which were more plastic in a familiar and unfamiliar context were also more unpredictable in these behaviours as well. There may be several reasons for this plasticity-predictability association. Firstly individuals which were more responsive to changes associated with repeated testing (i.e. were more plastic) may have also been more responsive to unmeasured stochastic fluctuations in their environment, making them more unpredictable (Bell and Sih 2008; Stamps and Groothuis 2010; Stamps et al. 2012; Bierbach et al. 2017). Secondly, there may be a genetic linkage between the genes controlling plasticity and the genes controlling predictability, which may be context dependent and sensitive to changes in the early development environment (Scheiner et al. 1991; DeWitt 1998; DeWitt et al. 1998; Charmantier and Garant 2005). This may be why correlations between plasticity and predictability were observed across both feeding treatments in an unfamiliar context but were restricted to only tadpoles in the high feed treatment in a familiar context.

Considering how individuals adjusted their behaviour across contexts, we only found evidence for a predictability syndrome in the high feed tadpoles. Thus, when developing under high feed conditions, individuals may be able to develop the same level of behaviour predictability across a range of contexts, whilst this may become decoupled under low feed conditions. This may be because behavioural predictability is energetically costly and can only be sustained across contexts by individuals with sufficient resources (Kwek et al. 2021; Mitchell et al. 2021b). It is also important to highlight that although we measured the total distance tadpoles swam in two contexts which differed in their novelty, we did not identify behavioural or plasticity syndromes between the two contexts under either of the development environments. This suggests that how an individual behaves initially in a familiar context and how they adjust their behaviour over time has no relation to how the same individual will behave in an unfamiliar context. Consequently, recording an individual’s exploratory behaviour in an unfamiliar context (commonly referred to as an open field trial) should not be used as a substitute for measuring their general levels of activity in a familiar context (Werner 1992; Werner and Anholt 1993; Anholt and Werner 1995; Babbitt 2001; Relyea 2005; Smith and Doupnik 2005; Royauté and Dochtermann 2017; Royauté et al. 2019).

At the residual level, we found that tadpoles which moved more for a given trial in a familiar context, subsequently moved less in the corresponding trial in the unfamiliar context or vice versa. As we only observed this correlation in the low feed treatment, this may reflect an energetic trade-off which low feed but not high feed tadpoles had to make. For example, low feed tadpoles may have had to manage their limited energy budgets so that if individuals travelled a further distance in a familiar context, they may have depleted their energy resources and travelled less distance within the unfamiliar context (Mathot and Dingemanse 2015; Peña-Villalobos et al. 2020). This result also suggests that the increased activity in the low feed treatment does not provide more resources to tadpoles since it would have led to the opposite relation. In contrast, high feed tadpoles do not show a negative correlation and suggest that they would have had the resources to invest an equal amount of energy in both contexts.

## Conclusion

By using a split brood design to control for genetic differences between individuals, we have shown that exposure to different early developmental conditions can trigger substantial changes in population and among individual variation in behaviour. Furthermore, how the behaviour of an individual is correlated across contexts is also highly dependent on the environment they experience during early development. Our results highlight the importance of early life effects on the development of among individual variation in behaviour and that early developmental conditions may only impact the phenotype at specific phenotypic levels and is highly context specific. Consequently, future research investigating how ecologically relevant development conditions may impact among individual variation in behaviour should incorporate multiple hierarchical levels of variance and investigate how an individual responds across multiple contexts.

## Author contributions

CB conceived the study, designed the experiment, performed the laboratory work, conducted the statistical analysis and wrote the first draft of the manuscript. PW and NC contributed to the study design and JM supervised the statistical analysis. All authors commented on the manuscript.

## Data Availability Statement

Data and analysis scripts are available from CB’s GitHub repository https://github.com/cammybeyts/xenopus_tadpole_food_availability. The code for the tadpole tracking tool has been provided by CB and can be accessed via GitHub https://github.com/cammybeyts/Tadpole_tracker.

## Acknowledgements

We would like to thank Mike Dye for his valuable help in assisting with laboratory preparations and tadpole husbandry and Craig Watt for building the flow-through system.

